# Organic electrolytic photocapacitors for stimulation of the mouse somatosensory cortex

**DOI:** 10.1101/2021.10.20.465090

**Authors:** Florian Missey, Boris Botzanowski, Ludovico Migliaccio, Emma Acerbo, Eric Daniel Głowacki, Adam Williamson

## Abstract

**Objective:** For decades electrical stimulation has been used in neuroscience to investigate brain networks and been deployed clinically as a mode of therapy. Classically, all methods of electrical stimulation require implanted electrodes to be connected in some manner to an apparatus which provides power for the stimulation itself.

**Approach:** We show the use of novel organic electronic devices, specifically organic electrolytic photocapacitors (OEPCs), which can be activated when illuminated with deep-red wavelengths of light and correspondingly do not require connections with external wires or power supplies when implanted at various depths *in vivo*.

**Main results:** We stimulated cortical brain tissue of mice with devices implanted subcutaneously, as well as beneath both the skin and skull to demonstrate a wireless stimulation of the whisker motor cortex. Devices induced both a behavior response (whisker movement) and a sensory response in the corresponding sensory cortex. Additionally, we showed that coating OEPCs with a thin layer of a conducting polymer formulation (PEDOT:PSS) significantly increases their charge storage capacity, and can be used to further optimize the applied photoelectrical stimulation.

**Significance:** Overall, this new technology can provide an on-demand electrical stimulation by simply using an OEPC and a deep-red wavelength illumination. Wires and interconnects to provide power to implanted neurostimulation electrodes are often problematic in freely-moving animal research and with implanted electrodes for long-term therapy in patients. Our wireless brain stimulation opens new perspectives for wireless electrical stimulation for applications in fundamental neurostimulation and in chronic therapy.

## 1. Introduction

Neuromodulation using electrical stimulation is a technique to activate or inhibit excitable tissues such as the brain directly via the delivery of an electric current. Electrical stimulation techniques were pioneered already in the beginning of the 20^th^ century to reveal function and connectivity of brain regions. One of the earliest notable achievements using electrical stimulation of the brain was the mapping of motor and sensory cortices by Wilder G. Penfield in 1937 (Penfield and Boldrey, 1937).[1] As a physician, Penfield further developed the electrical stimulation method known as the “Montréal Procedure”,[2] which is still employed today during resective surgeries in epilepsy where electrical stimulation allows the delineation of healthy brain tissue versus epileptogenic areas to be resected. Neuromodulation using electrical stimulation is now a standard tool in neuroscience, used to answer investigative questions in animal models and clinically established to improve outcomes of surgeries, distinguish the extent of brain tumors, [3] reducing adverse and unwanted disease effects in patients, and to treat various brain pathologies.[4,5] However, one feature common to all methods of electrical stimulation of brain tissue is the necessity to provide power to implanted electrodes via some manner of wire interfacing, typically via cables to power supplies or batteries. Batteries *in vivo* always have limited lifetimes and are restrictively bulky for convenient deployment in experimental animals.[6,7]

The established option in neuromodulation which does not involve the physical implantation of wires and batteries comes in the form of transcutaneous stimulation from electrodes located on or above the scalp, most commonly transcranial direct current stimulation (tDCS) and transcranial magnetic stimulation (TMS). An emerging approach is transcranial focused ultrasound stimulation, which can reach deeper structures with higher precision.[8] In 2005, Hummel et al. published one of the first studies using tDCS between two scalp electrodes demonstrating that tDCS can improve hand mobility in patients recovering from stroke. [9] Since then, many studies have shown that motor skills can be improved by noninvasive cortical stimulation via tDCS and TMS.[9–13] However, the main issue with transcranial methods is that they are almost exclusively used in hospitals due to, in the case of TMS, the complexity of the equipment and, in the case of tDCS, electrodes do not remain on the scalp permanently.[14] Recently, alternative modes of wireless stimulation using more minimalistic implantable devices have emerged. Approaches relying on power transmission via focused ultrasound,[15] magnetic field,[16] and light [17,18] all hold promise to resolve issues with wires and batteries for implanted devices.

Here, we explore the use of ultrathin, wireless, light-driven cortical stimulation electrodes. To this end we have adapted the recently-reported technology of the organic electrolytic photocapacitor (OEPC) [19]. The OEPC is activated when illuminated with impulses of deep-red wavelengths of light and correspondingly does not require connections with external wires or power supplies when implanted in tissue. Light is absorbed by an organic PN junction layer. In this work, we experimentally verify that the OEPC can stimulate the cortex directly in a wireless manner by converting light pulses which pass through skin and bone. The OEPC transduces illumination pulses into corresponding biphasic cathodic-leading electrical stimulation pulses. The OEPC was first reported in 2018 [20] for the stimulation of retinal explants *in vitro*, and has since been used in other *in vitro* electrophysiology studies.[19] Recently, OEPCs were successfully deployed *in vivo* for the stimulation of peripheral nerves.[21] Use of OEPC technology in the CNS has not been reported to-date. In this work, we apply ultrathin OEPCs directly onto the cortical surface. As a validation of efficacy, in this study we use the OEPC to activate the whisker motor cortex in mice and evoke behavioral and electrophysiological responses in the sensory cortex. Additionally, we demonstrate that the device can be coated with a thin layer of PEDOT:PSS, greatly increasing charge storage capacity of the device.[22] Importantly, the device can be actuated on the cortex using light transmitted through the skull and skin interface.

## 2. Materials and Methods

To enable moderately-invasive implantation of our wireless stimulation devices, we fabricated ultrathin and flexible OEPCs. **Figure 1** summarizes the general experimental procedure. OEPCs are fabricated using micropatterning techniques onto peelable parylene-C foils (see *experimental methods*). These devices feature arrays of PN sizes ranging from 1-3 mm ø. **Figure 1A**, shows devices on a glass wafer being peeled prior to implantation from the substrate using a drop of water. Details on devices fabrication and physics of operation have been covered in earlier studies describing their deployment *in vitro*. Specific experimental fabrication details can be found in the experimental methods section. Stimulation in this specific set of experiments is performed using a 638 nm laser, allowing the implanted device to be activated below the skin and skull without the necessity of a wire or tether to interface and power the electrical stimulus.

**Figure 1.**
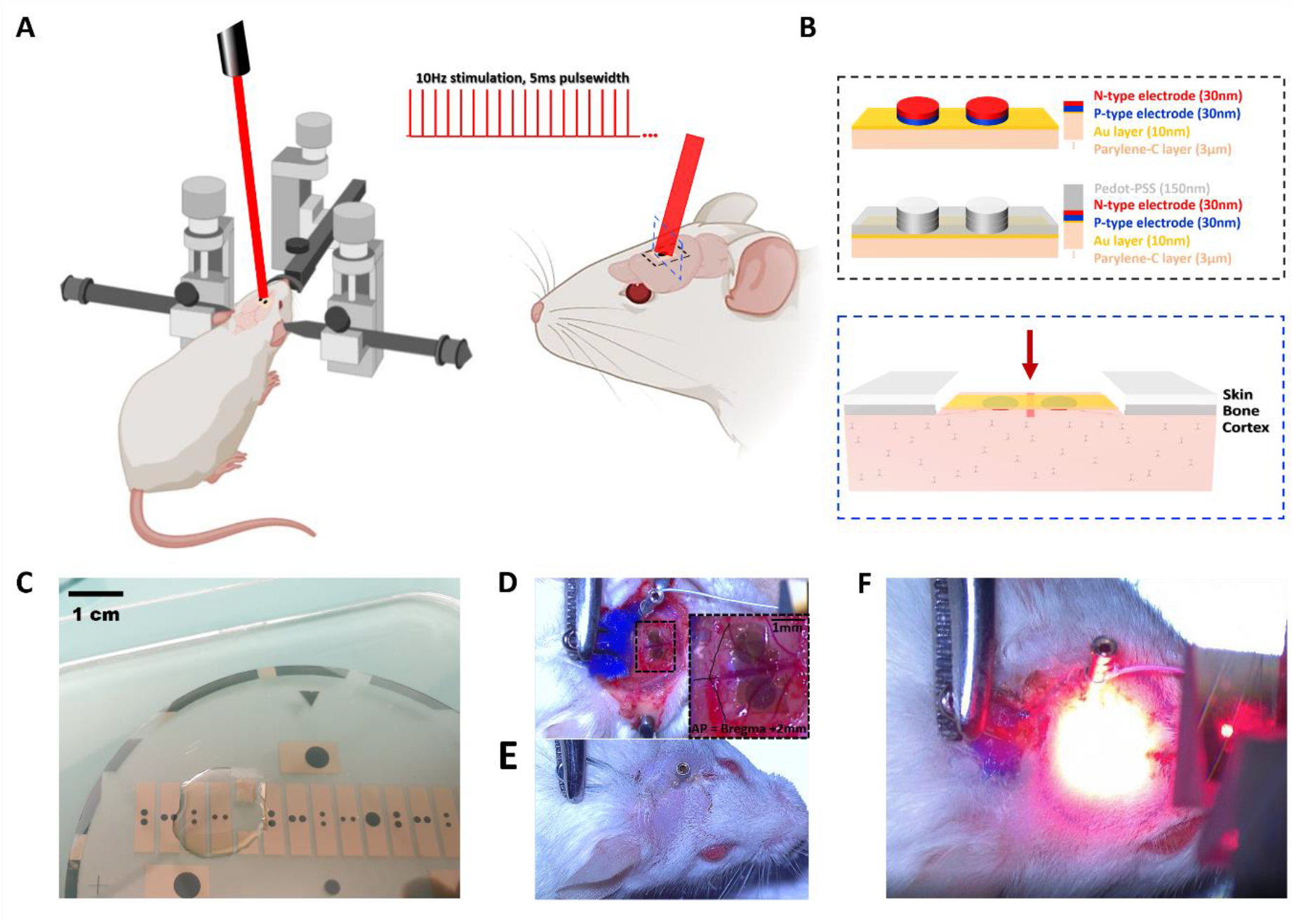
Organic photocapacitor implantation and stimulation on the mouse cortical surface. The experimental procedure is summarized in the illustrations and photographs shown here. **(A)** Anesthetized mice are placed in a stereotactic frame, a craniotomy opens the skin and skull depending on the desired configuration. A laser diode (638 nm) mounted above the stereotax is used to deliver stimulation impulses (5 ms duration, 10 Hz) to the implant site. Devices are tested acutely. **(B)** Cross-section of the flexible OEPC stimulator. The device features a PN junction acting as a charge-generating electrode, and an underlying semitransparent return electrode. Variants with and without a PEDOT:PSS coating are used in this work. The OEPC device is laminated on top of the exposed cortex, and light pulses are delivered from above. **(C)** OEPC devices are fabricated on parylene-c substrates which are lithographically-defined on a glass substrate. This image shows delamination of an OEPC device from the glass wafer immediately prior to device implantation. Water is used to lift-off the parylene-c from the underlying hydrophilic glass. **(D)** OEPC device implantation on the surface of the mouse cortex above barrel motor areas in both hemispheres to evoke whisker movement. Stimulation is performed using three illumination configurations: Directly on the cortex **(E)**, illuminating through bone + cortex (not shown), and through skin + bone + cortex **(F)**. Skin and bone are relatively transmissive to the 638 nm wavelength light, therefore the OEPC can be activated when fully implanted under skin+bone. Stimulation artefacts and evoked sensory response were recorded using metal electrodes on the primary sensory (S1) cortex with behavioral whisker movements recorded simultaneously using video.

### 2.1. Flexible Organic Electrolytic Photocapacitor (OEPC) fabrication

The active organic PN materials [H2Pc, (Phthalocyanine, Alfa Aesar) and PTCDI (*N,N*′-dimethyl-3,4,9,10-perylenetetracarboxylic diimide, BASF)] are prepared and deposited the way as described in previous studies [19–20], however fabrication and patterning details are different in this work. 4-inch soda lime glass wafers (University Wafer, 550 μm thick) were cleaned in a 2% solution of Hellmanex III detergent heated to 50°C for 30 min, followed by a high-pressure rinse with acetone and deionized water (DI). Substrates were then treated with O_2_ plasma (Diener GmbH, 200 W, 20 min). Immediately after, parylene-C (5 μm thickness) was deposited via chemical vapor deposition (Diener GmbH). The parylene-C was subsequently patterned via lithography and reactive ion etching (RIE) to produce 4 ×15 mm ribbons. Patterning was done as follows: 80 nm of Al was evaporated through a shadow mask to define the areas of the parylene ribbons. The surrounding unmasked parylene was removed by RIE (200W, O_2_ 100 sccm). The Al mask was then etched using a commercial wet etch solution. The surface of the parylene was then activated using O_2_ plasma (50 W, 2 min), followed by vapor-phase treatment with (3-mercaptopropyl)trimethoxysilane, MPTMS, by placing the samples in a chamber saturated with MPTMS-vapor heated to 90 °C for 1 h. MPTMS treatment improves the adhesion of Au on the parylene-C substrate. Next, a semitransparent 10 nm-thick film of Au was thermally evaporated over the whole wafer in a vacuum of < 2×10^−6^ Torr using a rate of 3-5 Å/s. The organic pigment PN pixels were formed by thermal evaporation through steel shadow mask (75 micron thickness) at a base pressure of < 2 × 10^−6^ Torr using a rate of 0.1-0.5 nm/s. 30 nm of H_2_Pc and 30 nm of N-type PTCDI were successively deposited resulting in 60 nm total thickness of the organic layers (PN). PEDOT:PSS modification of the OEPC device was performed as-follows: Clevios PH1000 PEDOT:PSS + 2% w/w (3-glycidyloxypropyl)trimethoxysilane (GOPS) was spin-coated at 1500 rpm using a 1000 rpm s^−1^ acceleration for 60 s to obtain a well-adhered PEDOT:PSS coating (150 nm ± 5 nm, measured using scanning stylus profilometry). The films were subsequently baked at 130 °C for 1 h and were immersed in 0.1 M KCl to remove any excess low-molecular weight compounds as well as to allow the PEDOT:PSS to take up water and swell. Samples were then stored in 0.1 m KCl for 20 h before further use.

### 2.2. Animals

Animal experiments were done with the agreement of European Council Directive EU2010/63, and French Ethics approval (Williamson, n. APAFIS 20359 – 2019041816357133 accepted by comité d’éthique en expérimentation animale n°071). Experiments were performed on 12 OF1 (Oncins France 1, from Charles Rivers Laboratories, France). These OF1 mice were adult mice from 8 to 10 weeks old. Mice were submitted to a normal cycle of 12/12h day-night cycle and kept at room temperature. The animals were housed in transparent cages in a group of four, water and food were dispensed *ad libitum*. A total of 12 mice were used in this study.

### 2.3. Surgical procedure

Mice underwent anesthesia with a xylazine (20mg/kg) and ketamine (50mg/kg) mix with intraperitoneal injection of 2.5 μL/g of the mix. During the experiment, the experimenter made sure that mice eyes were covered with vitamin A. After a subcutaneous lidocaine injection, a small skin incision was made on the skull where it had been previously shaved. A modified metal clip was used as an anchor point on the skull. The clip was glued to the skull using a dental cement primer (Ultradent®) followed by a resin photopolymerized by an UV lamp (Lc Block-Out Resin from GACD®). The clip was then anchored to an articulated arm (Noga®, RS-Components) to stabilize the mouse skull. Once the skull was tightly fixed, a craniotomy using a Dremel (K1070 from Phymep®) and 0.2 mm drill bit was realized on top of the barrel motor cortex (AP : +0.38 mm; windows of about 4 × 1.5 mm). *Dura Mater* was removed and the OEPC pair was gently deposited on the whisker motor cortex, each PN contact on a different hemisphere. A small burr hole was drilled next to the craniotomy (AP = Bregma −0.9mm; ML = Bregma – 3mm) with a 0.8 mm drill bit and *Dura Mater* was also removed. A mini pin (RS Component®) was inserted until it was in contact with primary sensory cortex, the pin will act as a recording electrode to monitor stimulation artifacts. It was then cemented similarly to the clip was secured. During light stimulation sessions, the eyes of the mouse were covered by aluminum foil to prevent any spurious stimulation via the visual system.

### 2.4. Light stimulation and recording

Continuous brain recording was obtained with an Intan 128ch Stimulation/Recording Controller (IntanTech®). mice were lightly stimulated using a One Channel LED Driver 10A associated with a 638nm Diode Laser. To aim the laser beam toward the photocapacitor location, the laser power was set at 5% and in continuous mode. Under visual examination the laser beam was then shifted until it reached the OEPCs. Once the laser beam was at the right location, the power was set at 100% (700 mW, max current 1000 mA) and the stimulation protocol started simultaneously with the brain recording. Each stimulation session was composed of 150 stimulations with a 5 milliseconds ON-time and 95 milliseconds OFF-time to reach a total 15-second-long stimulation. All stimulation sessions were spaced by 5 minutes to let the brain rest. In order to reduce the number of used animal and increase the statistical power of the study, all mice underwent the three protocols (exposed cortex stimulation, stimulation with bone and stimulation with bone and skin).

### 2.5. Computational simulation methods

All finite elements simulations were made using COMSOL Multiphysics® software, version 5.5. A simpler brain mesh was sculpted using Paxinos mouse brain as a model. A disc representing the photocapacitor was then placed above the target area. All the meshes were scaled appropriately, in order to fit the *in vivo* experiment. Frequency dependent physical properties like permittivity and conductivity were applied to all the components in the model (see Figure S1). The results of the simulation come from the ‘electrical current’ study. Stimulation voltage value in the finite element simulation study was set to the same value delivered by the photocapacitor stimulation experimentally recorded in Figure 2. Electric field lines and electric potentials were plotted in the 3D brain to illustrate and characterize the effects of the *in vivo* stimulation.

**Figure 2.**
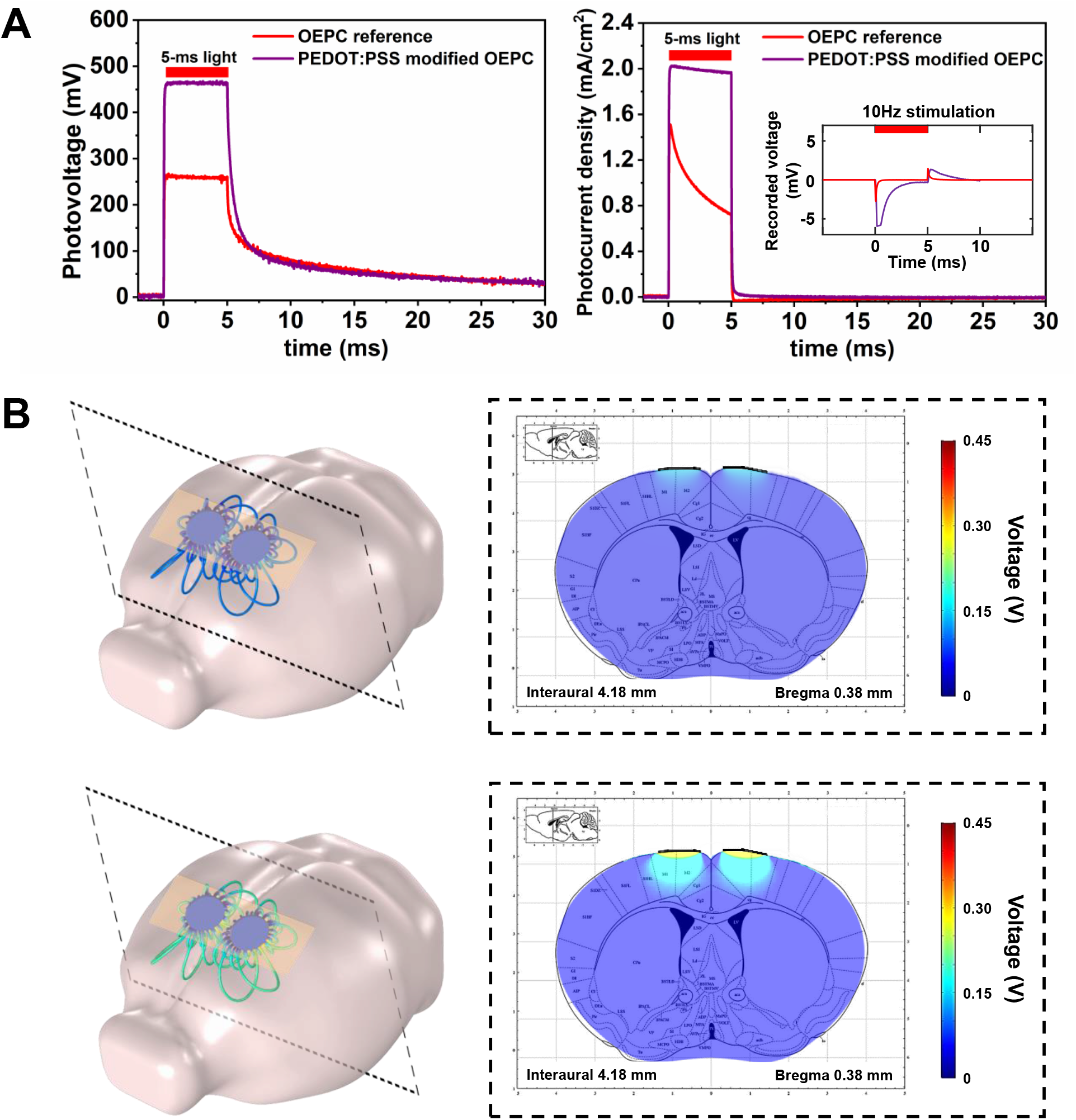
OEPC photoelectrical characterization and modeling of stimulation. (**A**) Both standard OEPCs (in red) and PEDOT coated OEPCs (in purple) were tested *in vitro* by measuring electrophotoresponse (EPR). EPR measurements are conducted by measuring photovoltage and photocurrent between the bottom contact (gold) and an electrolytic contact confined on top of the PN region (Ag/AgCl in 0.1M KCl solution). This configuration measures only the cathodic phase of the photocharging. Devices are illuminated from the bottom, that is through the gold layer, using a 638 nm laser diode with 5 ms pulses at a 33 mW/cm^2^ optical power. PEDOT-modified devices show substantially better photovoltage and photocurrent values than unmodified analogs. *Inset*: Transient voltage stimulation artefact recorded in an agarose gel brain phantom, using a recording electrode placed in the gel near the center of the OEPC pixel, versus a distant ground electrode. This transient voltage measurement clearly resolves the biphasic voltage perturbation produced by the OEPC upon pulsed illumination. This voltage transient is recorded using 5 ms light pulses with 10 Hz repetition. **(B)** The photovoltage peak values were subsequently implemented in our finite element model (FEM) of mouse brain in Comsol Multiphysics® as the maximum voltage difference between two PN photoelectrodes and a gold return electrode. FEM allows estimation of the potential perturbations that can be induced by the device in the brain. A coronal slice taken from the simulations indicates an increased potential penetrating the cortex for the PEDOT:PSS covered devices.

### 2.6. Signal processing and statistical analysis

Previous to experiment, we calculated with a power analysis calculator the number of animal needed when using paired samples and a global power of 0.8. Matlab was used to analyze row Local Field Potentials (LFP) as well as filtering data. LFP recordings were collected with a sample rate of 30K samples per seconds and a 50 Hz notch filter was directely applied. In addition of the 50 Hz notch filter, a 10 Hz low-pass filter was applied in Matlab. Both Figures 3 and 4 represent examples of filtered recordings acquired respectivelly during the light stimulation and right after the stimulation. To perform this analysis, we segmented our recordings in groups of 10 seconds recordings for ‘during stimulation’ and ‘ right after stimulation’. Once all the recording were semented in these two groups we gathered them regarding the type of OEPC and the stimulation protocol used. In order to visualize the effect of the stimulation on the LFP recordings, time/frequency analysis were performed on post-stimulation recording to correlate the behavioral result of the stimulation with the increase of activity in the time/frequency plot. Regarding stimulation artefact, raw recording during the stimulation were plotted and the average absolute value of the stimulation artefact was calculated over the 10 seconds of stimulation. Data were collected with this method for each of the 12 mice (2 groups of 6 mice) and combined into an Excel file to perform statistical analysis For the statistical analysis, we used the software R. Before any statistics, we performed normal distribution tests (Shapiro tests) to adapt our analysis regarding the normality of the data. Pairwise-wilcoxon tests (α = 1%) were conducted and p-values were calculated to highlight some potential significant differences in the data. P-values were reported on the boxplots plotted in Figure 3 and 5 to better illustrate the differences between the different groups. Finally, plots were made and analyzed using both Matlab and R.

**Figure 3.**
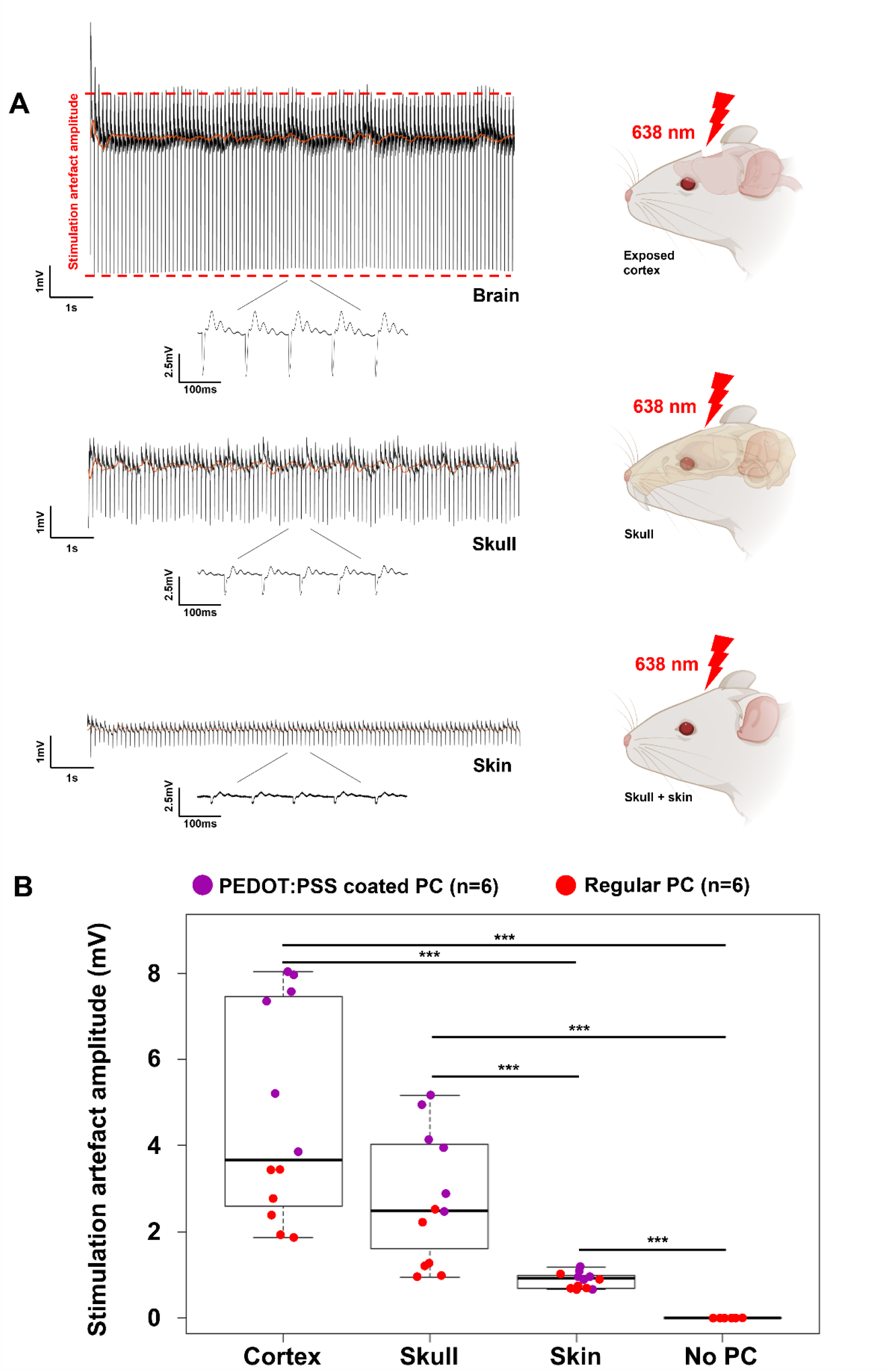
Characterization of stimulation artefacts amplitudes for the 3 protocols. Raw stimulation artefact recordings are shown in **A** to characterize the stimulation amplitude delivered by the OEPCs under conditions with illuminating through the bone or bone+skin. Evoked biphasic pulses are represented as insets for all three stimulation protocols. (**B**) The recorded stimulation artefact amplitude when illuminating the device directly on the cortex, or with intervening skull and skin+skull (p-value < 0.001). Regarding the effect of PEDOT:PSS coating on stimulation artefact amplitude, we could find significant differences when looking at the Cortex and Skull groups (p-value_cortex_ = 0.0022, p-value_Skull_ = 0.0043) but no difference was highlighted in the Skin group (p-value_skin_ = 0.3701).

**Figure 4.**
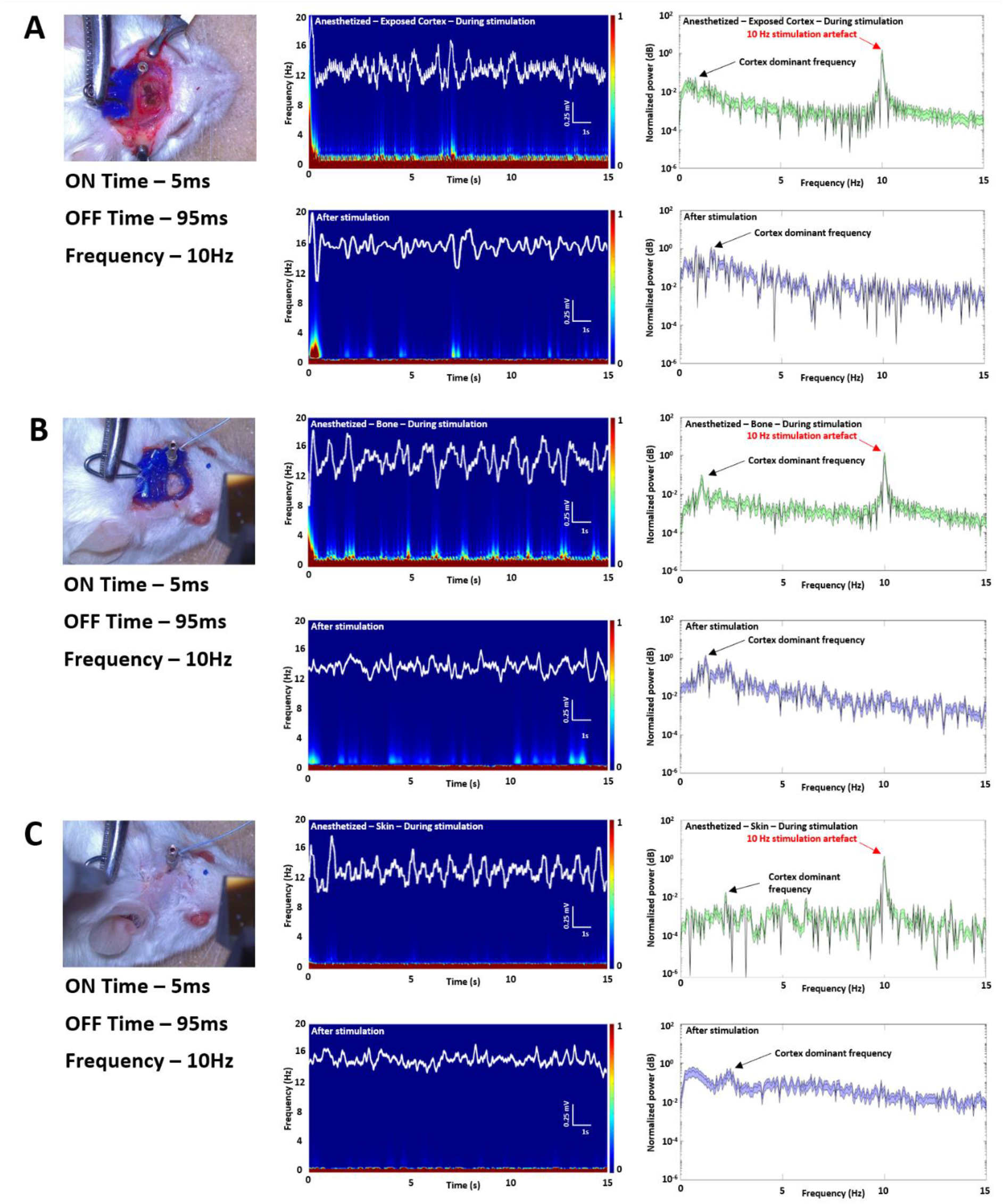
*In vivo* stimulation of the motor cortex in anesthetized mice. Each recording has been analyzed using a low pass 10 Hz filter to sort real biological activity from the stimulation artefact noise. 3 stimulation protocols were performed **A)** OEPC on the exposed cortex (n = 12 mice, dispatched into two groups of 6 mice for regular OEPCs or Pedot:PSS coated OEPCs implantation), **B)** OEPC covered with bone alone (n = 12 mice with 2×6 mice) or **C)** OEPC covered with bone and skin (n = 12 mice with 2×6 mice). For all three stimulation protocols, LFP were recorded in the somatosensory cortex (solid white line in the Time/Frequency insets) and Time/Frequency analysis was performed. All recordings have been segmented and a power spectrum analysis was done on all the segments showing a clear 10Hz artefact during the stimulations protocol and the intrinsic cortex frequency around 2 Hz for the anesthetized mice (more difficult to see in A for non-ideal open craniotomy recordings, very clear in B and C with closed craniotomy recordings). Clear activation can be seen during the stimulation. That activation correlates with whisker movement in behavioral analysis and stops as soon as the whiskering stops.

**Figure 5.**
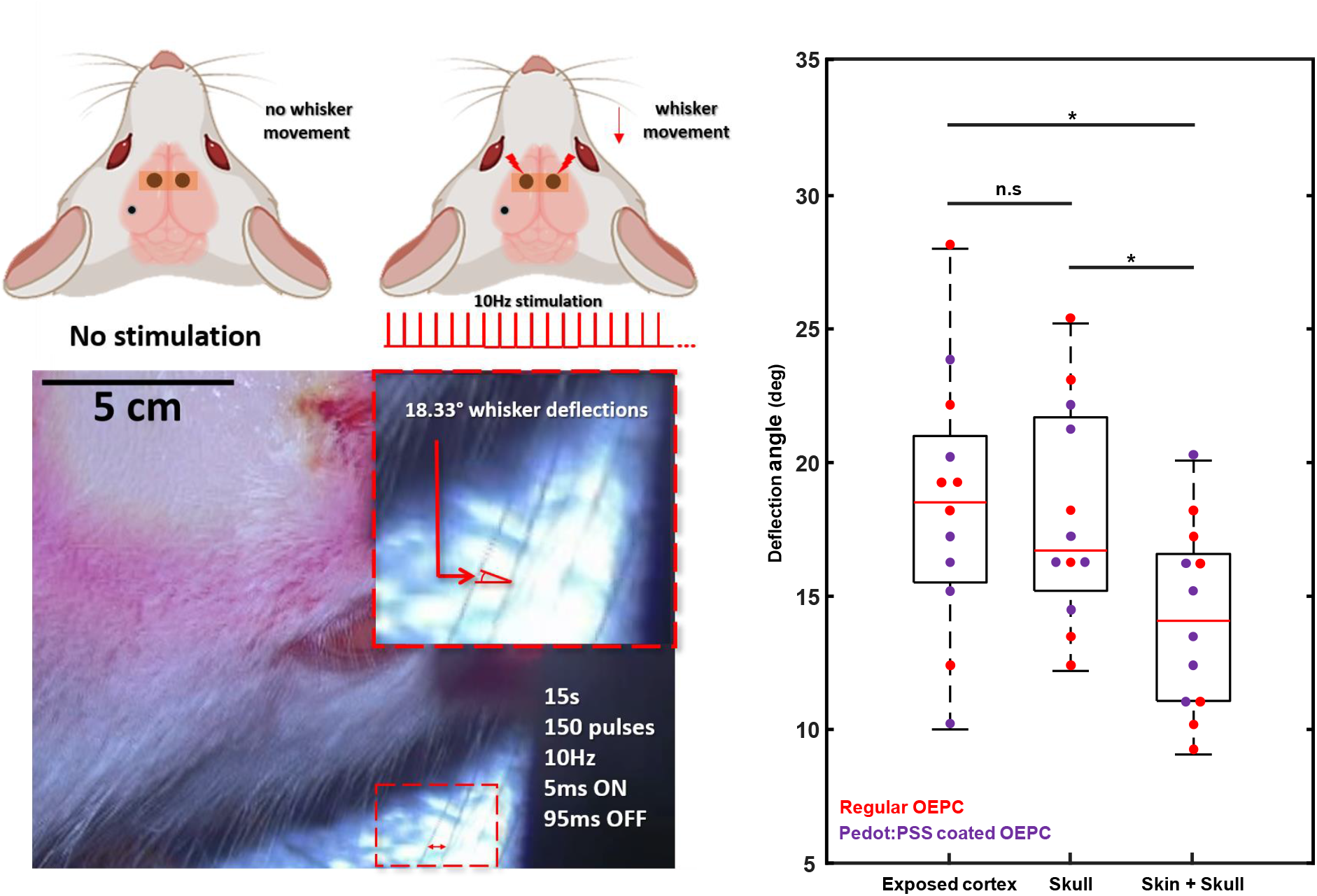
Whiskering evoked via OEPCs. Stimulation applied to the cortex, with light on exposed cortex, through the skull or penetrating the skin and skull to activate the OEPC on 12 mice divided into 2 groups of 6 for regular or coated OEPCs. As plotted and shown in the overlaid-video image, 10 Hz light pulses evoke an average whisker deflection of 18.33° during stimulation which corresponds well to literature where activation of the somatosensory region in mice with 5 Hz to 15 Hz gives a maximum whisker deflection of 28°.

## 3. Results

### 3.1. *In vitro* characterization of OEPC electrophotoresponse (EPR)

OEPCs convert impulses of light into biphasic current pulses: a cathodic one at the start of the light pulse, and an anodic one once the light pulse finishes.[23] OEPCs are characterized prior to implantation using an experimental setup for measuring EPR described in previous literature.[24] EPR registers the photovoltage and photocurrent which is produced by the OEPC device upon pulsed illumination in contact with electrolytic solution. When measuring between the back contact and an electrolytic contact of Ag/AgCl in 0.1M KCl, the EPR method quantifies the charging photovoltage and photocurrent of the cathodic phase only. The measured photocurrent provides a quantitative expectation of stimulation current values which are injected into tissue. The EPR traces for the OEPC devices are plotted in **Figure 2A**. OEPCs illuminated through the semitransparent gold bottom electrode with 33 mW/cm^2^ optical power produce peak electrolytic currents of around 1.5 mA/cm^2^. Over a 5 ms light pulse, a charge injection capacity of 7.48 μC/ cm^2^ ± 0.21 (n=4) is measured. Based on recent findings [25] on rigid OEPC devices on indium tin oxide glass, modification of the OEPC electrode surface with a thin layer of the conducting polymer formulation PEDOT:PSS leads to decreased electrode impedance and higher charge injection capacity. With the flexible parylene-C/Au OEPCs tested in our work, we found that PEDOT:PSS modification (150 nm thickness) leads to substantial improvements in performance. Due to the substantial increase in charge capacity, photocurrents of 2 mA/cm^2^ are sustained over a 5 ms illumination pulse. This corresponds to a cathodic phase charge injection of 10.01 μC/ cm^2^ ± 0.27 (n=4) that can be expected for a 5 ms light pulse. These photocurrents and charge densities (using the same illumination conditions) are over twice as high as previously-published best on rigid PEDOT-modified OEPCs [25], and significantly higher than other published PEDOT-based photostimulation devices.[26] In parallel to the EPR method, we characterized also the OEPC in electrically “floating mode”, recording the voltage transients produced in solution upon illumination at a 10 Hz frequency (**Figure 2A, inset**). The biphasic nature of the stimulation transient is clearly visible in this case, as is the substantially higher amplitude of PEDOT-modified devices versus unmodified controls. Both types of OEPCs easily transduce stimulation frequencies up to 100 Hz (Figure S2). For stimulation of the mouse somatosensory cortex, we proceeded to test the two types of OEPCs, both with or without the addition of a 150 nm layer of PEDOT:PSS on top of the device. To achieve stimulation of both motor whisker areas to evoke whisker movement on both sides of the animal, we employed a dual-pixel configuration. **Figure 2B** takes the values of photovoltage and includes them in a finite element model of the mouse brain. The devices are modeled over the barrel region of the somatosensory cortex, field vales can be seen to penetrate deeper into the cortex as expected for the devices covered in the additional PEDOT layer.

### 3.2. OEPCs wirelessly stimulate the whisker motor cortex and induce electrophysiological activity

For the photostimulation protocol, we used a 638 nm diode laser, giving 700 mW of optical power through beam shaping optics to produce a 20 mm^2^ illuminated area. Therefore, a 5 ms length pulse yields 17.5 mJ/cm^2^. This light dose is more than ten times below the laser safety standards for skin exposure according to the *American National Safety Standard for the use of Lasers,* according to which maximum permissible exposure at 638 nm is 195 mJ/cm^2^. Experiments were performed in the configuration as shown in **Figures 1B**: anesthetized mice were placed in a stereotaxic frame and a craniotomy was performed to expose the skull or cortex surface. For all *in vivo* experiments, double OEPCs using the PEDOT:PSS coating were implanted to the target region of the cortical surface. In addition to the double OEPCs placed on top of both whisker motor areas, a metal recording electrode was placed in the barrel sensory cortex to record evoked activity as well as stimulation artefacts. The recording electrode position is 3 mm deep into the brain, away from the OEPC surface. Recording stimulation artefacts allowed relative quantification of stimulation amplitude (**Figure 3**).

The recorded stimulation artefact scales with the three protocols of photostimulating directly on the bare cortex, through bone, and through skin and bone. Thicker intervening tissue, not surprisingly, leads to lower stimulation amplitude. PEDOT-modified devices performed better than unmodified ones. It is important to note that control photostimulation of the cortex without an OEPC device yielded no measurable signal. In **Figure 4** we show the stimulation-evoked S1 response during stimulation and immediately after the stimulation for the three protocols (bare cortex, through bone, and through skin and bone). For our 3 protocols we can distinguish S1 cortex activation during the stimulation with a frequency around 7 Hz to 9 Hz, corresponding to an activation of whiskers in the M1 cortex.[27] This cortical activation correlates behaviorally with a global movement of all whiskers with no significant differences between the PEDOT:PSS coated OEPC group and the regular OEPC group (p-value_wilcoxon_ = 0.3723, **Figure 5**). As we replace tissue on top of the motor cortex, the activity in S1 becomes weaker and flatter but still distinguishable during the stimulation. After the end of the 15 s stimulation period, no more behavioral activity was seen on the whiskers and the 7 Hz −9 Hz signal was no longer present on the electrophysiological recordings. The decrease in S1 response can be directly explained by the intensity of the stimulation itself. When analyzing the amplitude of the recorded stimulation artefact (**Figure 3**), we observe a clear decrease in the amplitude of the recorded artifact with each additional layer of tissue in place. The whiskering behavioral response does not show significant differences between the different stimulation conditions, which is most likely because in the present OEPC device configuration, a relatively large area of the cortex is being targeted without high specificity.

## 4. Discussion

Electrical stimulation of brain tissue can be effectively provided by electrodes using both invasive or non-invasive techniques. Generally, non-invasive methods like transcranial magnetic,[28] electrical,[29] or focused ultrasound[8] stimulation are not very specific and also have a limited depth of activated targets. Implanted stimulation technologies carry the risks and complications of surgical procedures, however they can target structures more precisely and at-depth. To achieve superior implantable stimulators, the goal is to make devices wireless and as small and minimally-invasive as possible.[30,31] Various approaches relying on inductive power transfer,[32] magnetic resonant coupling, focused ultrasound, or optical stimulation are being actively explored at the level of *in vivo* animal experiments.[7] In our work, we have explored ultrathin photovoltaic-based implants for stimulation. Such microengineered optoelectronic devices can offer a solution to achieve wireless stimulation. Light is convenient since LED sources are available in a large variety and at low cost, and the use of light is established in medical practice. OEPC devices are a highly miniaturized flexible optoelectronic stimulation system. The devices used in this work are fabricated on parylene substrates with a total thickness of approximately 4 μm. OEPCs have, to-date, not been tested in the CNS.

Our experimental findings show that OEPC-mediated electrical stimulation is activated by a deep-red light source which penetrates tissue reaching implanted devices without the necessity of directly contact the device. The OEPC implants transduce impulses of light into electrical stimulation pulses, and achieve comparable results in terms of behavioral and electrophysiological response in *in vivo* experiments done using electrical stimulation or optogenetics.[27,33,34] Whisker movement after electrical or optogenetic stimulation has been well described in the literature [35] but the behavioral output of M1 electrical stimulation is not fully understood. Although M1 stimulation should result in a rhythmic protraction of the contralateral whiskers it has been demonstrated that a spreading long train of stimulation (especially to S1) can produce highly different behaviors.[33] This wide stimulation going from M1 and spreading to S1 can explain our complex behavioral results (rhythmic retraction of contralateral whiskers). It is worth the comment that optogenetics leverages the versatility of light, however the OEPC has two distinct advantages: 1) It does not necessitate any genetic manipulation and 2) OEPCs operate with deep red wavelengths which have relatively good tissue penetration. Opsins used in optogenetics typically are sensitive to blue and green wavelengths and very few are available with response to red light. We have demonstrated that the optoelectronic stimulation provided by the OEPC is directly correlated with the light power reaching it. Coupling the ability of deep-red light to easily pass through tissue and the property of OEPC to transduce light into stimulation current enables an increased flexibility in experimental protocols. We established that the deep-red light laser was powerful enough to elicit a physiological response through the fur, skin and skull combined. As some degree of light is lost passing through biological tissue, we additionally showed that PEDOT:PSS coating is one solution to allow a better efficiency of the OEPC (≃ doubling the charge injection capacity) when stimulating brain tissue.

## 5. Conclusion

Methods of brain stimulation generally fall into two categories, invasive or non-invasive. Regardless of the level of invasiveness, stimulators must necessarily be connected via wires to forms of power supplies to provide current to electrodes. Our work was motivated by finding new ways to accomplish wireless stimulation to enable experimental protocols which are difficult or impossible using wired/tethered systems. Light is certainly one of the less-explored modalities for wireless transcutaneous/transcranial stimulation, and our work shows that this approach has promise.

Our prototype OEPC stimulator is activated via externally applied deep-red light which penetrates skin and bone to reach the minimally-invasive devices located on the mouse cortex surface. In the work here, we activated the devices and characterized the degree of signal attenuation when increasing the depth of implant (beneath the skin, beneath the skin and bone) to demonstrate the ability of the red light to reach and activate the device. We modeled the results using finite element methods to similarly show the device and its estimated performance on the cortex surface.

Practically, we utilized the device as a minimally-invasive motor cortex stimulation device, specifically targeting the barrel cortex of mice. The stimulation demonstrated a well-controlled activation of whiskering and the deflection of whisker movement was characterized.

The OEPC stimulation technique would be an interesting addition to a wide range of freely-moving neuroscientific protocols which utilize electrical brain stimulation, but suffer from a complex arrangement of tethers to provide power to electrodes. Indeed, tethers and wires in classical electrical stimulation methods can disturb social interactions between animals and free exploration in animal cages, thus hindering normal behavior in behavioral studies.[36,37] From a device optimization point of view, successful deployment of OEPC implants requires increase of the charge density/light input ratio, in order to generate charge efficiently and safely. This would allow the use of this approach to reach deeper targets. Tissue transparency and safe optical light delivery is relatively well-documented and understood for skin, muscle, and fat tissues, in large part due to the prevalence of photodynamic therapy for cancer.[38] A recent report[21] showed successful OEPC stimulation of a peripheral nerve, actuated with light (100-1000 μs total optical pulse length) transmitted through 10-15 mm of tissue. This suggests that OEPCs can be used in transcranial stimulation in larger, thicker-skulled animals models, since the transmittance of wavelengths in the red-near infrared region is very similar for bone as it is for skin.[39] The present proof-of concept study validates stimulation of the cortex and the experimental principles of using such devices. Next steps should also focus on stimulating more specific areas of the cortex using devices with smaller stimulating electrodes, for instance ones designed to concentrate photogenerated charge on a smaller, selectively-exposed electrode. Such OEPCs would deliver more spatially-confined electrical stimulation with higher charge density, leading to more precise and effective behavioral responses. Additionally, the photogeneration efficiency will dictate the minimal optical pulse length needed for stimulation. This efficiency is also important for considering unwanted photothermal effects. Applications with high-frequency stimulation or high optical duty cycle can result in heating. For chronic applications, stability and biocompatibility are important. Parylene-C [40] is chosen due to good indications of bioocompatibility, however in principle other substrate materials can be easily substituted. The long-term biosafety of PEDOT appears promising,[22] while the PN organic materials are well-known industrial organic colorants established to be nontoxic an used in commercial products for many years.[41] The next step for testing OEPC efficacy will be chronic tests.

OEPC technology could potentially be translated to the clinic for on-demand moderately invasive stimulation in humans. The design, shape and size of these devices can be highly customized and scaled to accommodate patients, not simply small rodents, with clear applications for stimulating a wide variety of cortical zones. Future research directions should focus on improving these devices in terms of charge injection capacity and layouts of devices to stimulate deeper brain structures.

## Acknowledgments

This project has received funding from the European Research Council (ERC) under the European Union’s Horizon 2020 research and innovation program (A.W. grant agreement No. 716867; E.D.G. grant agreement No. 949191). E.D.G. and L.M. gratefully acknowledge financial support from the Knut and Alice Wallenberg Foundation within the framework of the Wallenberg Centre for Molecular Medicine at Linköping University and the Swedish Research Council (Vetenskapsrådet, 2018-04505).

## Author contributions

E.D.G. and A.W. conceived the project. B.B., and F.M. performed experiments. L.M designed and fabricated organic photocapacitor devices. F.M. performed finite-element simulations. F.M. analyzed neural data, A.W. wrote the paper with input from the other authors, including E.A and E.D.G.

## Supplementary information

**Figure S1.**
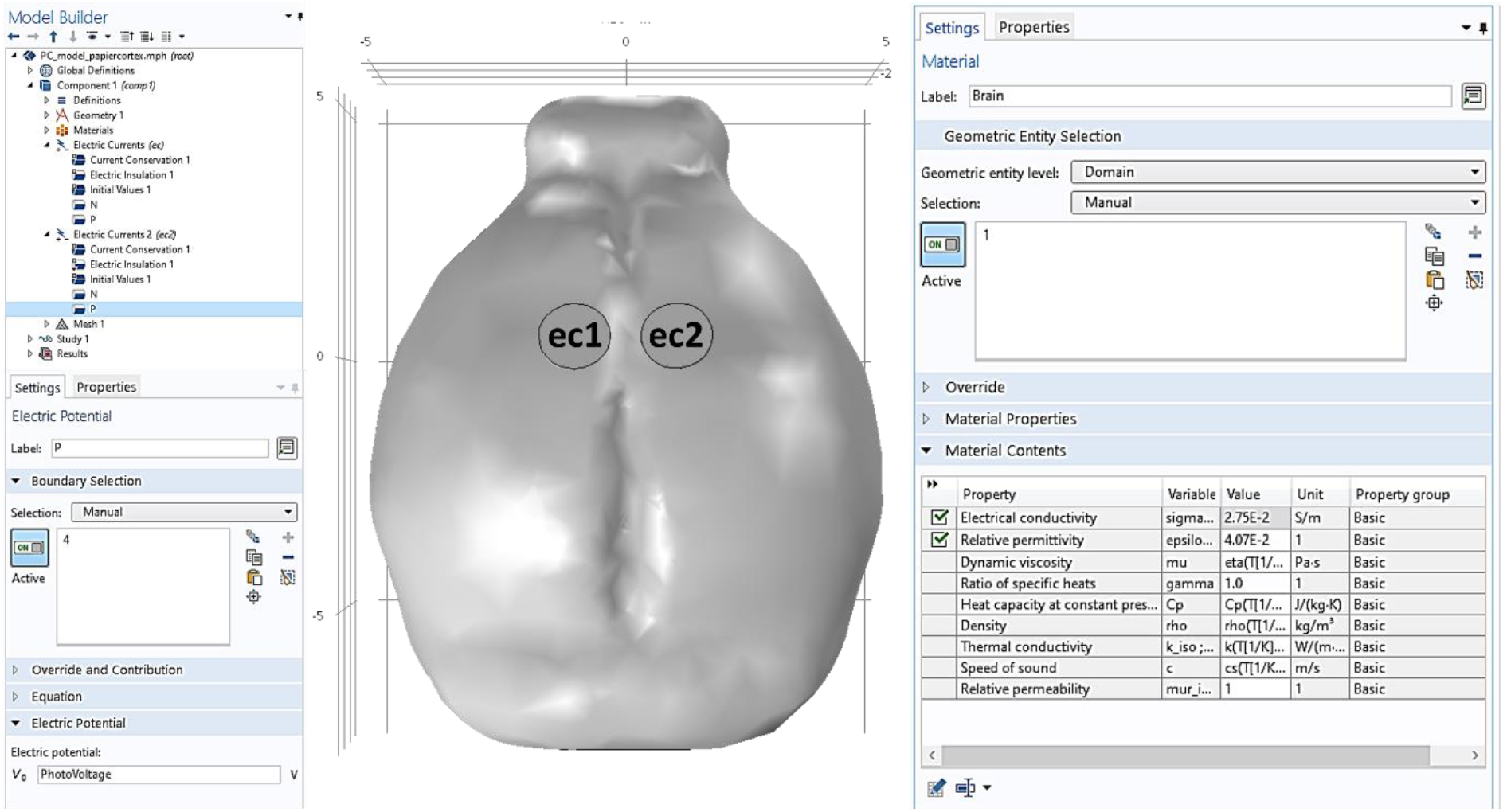
FEM parameters for PC stimulation modeling.

**Figure S2.**
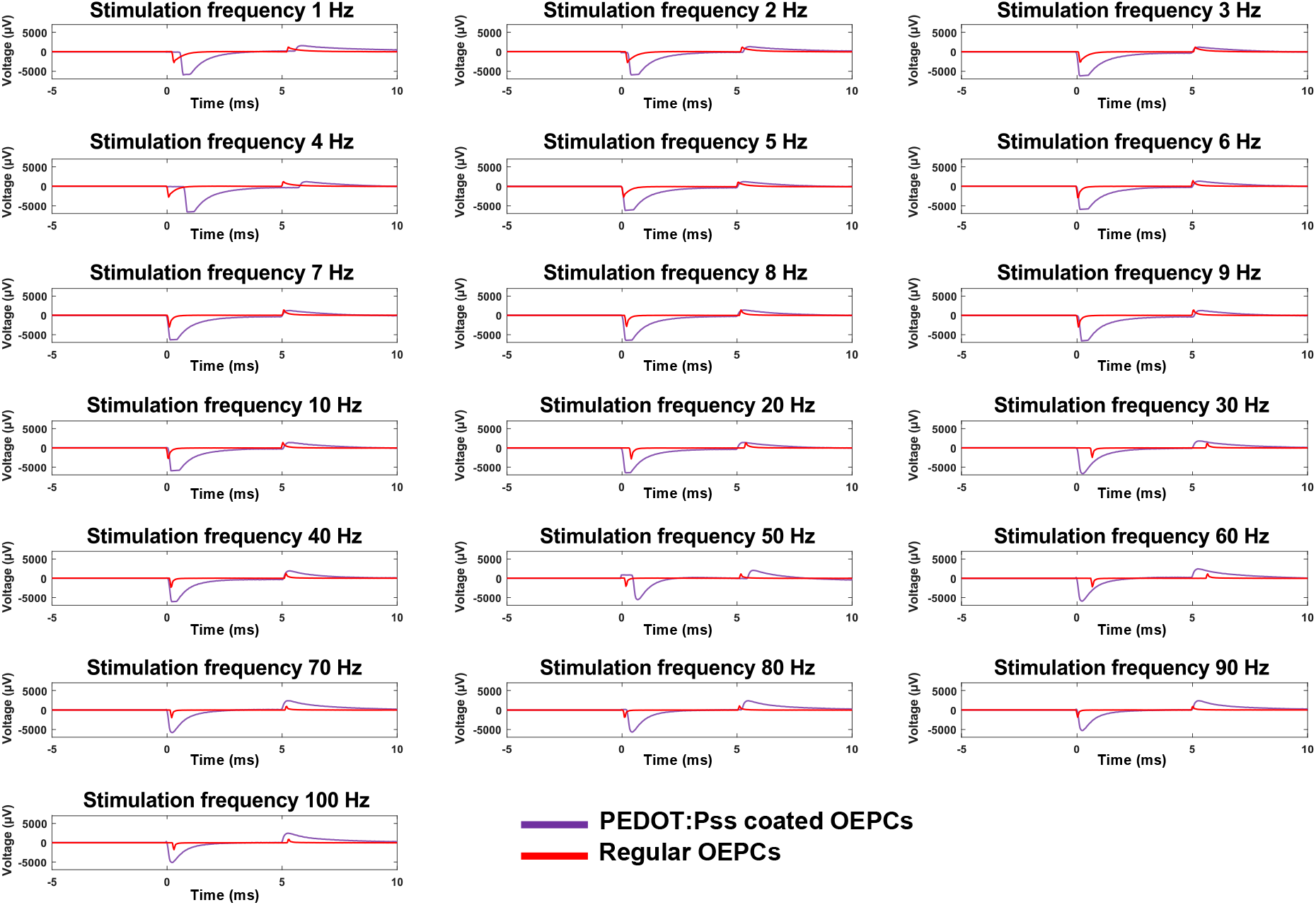
Recorded induced tranient voltages (averages of 20 consecutive pulses) during frequency-dependant pulse stimulation for regular and PEDOT:PSS coated OEPCs.

